# Attention reshapes the representational geometry of a perceptual feature space

**DOI:** 10.1101/2025.08.28.672962

**Authors:** Angus F. Chapman, Melissa Allouche, Rachel N. Denison

## Abstract

Department of Psychological and Brain Sciences, Boston University Our perception of the world is transformed by attention, both in terms of the efficiency of information processing and the appearance of attended stimuli. A standard theory is that attention regulates how competing stimuli vie for resources. However, an alternative perspective is that attention alters the representational geometry of stimulus spaces, such that changes in processing are not isolated to the particular competing stimuli, but are reflected across the entire perceptual space. To test this representational hypothesis, we conducted an experiment in which participants reported the perceived similarity of orientations spanning the full stimulus space when attention was directed or not directed to specific orientations. We used these similarity judgments to measure the representational geometry of orientation, finding that attention reliably expanded the representational space in a narrow range around the attended orientations. We also found evidence for compression of the representational space in a broad range around unattended orientations. Our findings support the idea that attention acts to reshape the representation of entire perceptual spaces in a way that supports processing of relevant stimulus features. By simultaneously manipulating attention while measuring perceptual similarity, our methodological framework opens the door for future work investigating the interaction between cognitive and perceptual processes from the perspective of representational geometry.

---

Attention shapes the way we perceive the world around us. We can direct our attention to spatial locations, enhancing our ability to detect or discriminate objects at those locations (Carrasco, 2011). Similarly, attending to relevant features of objects, such as their color or direction of motion, increases our sensitivity to those features. Findings from a broad range of studies have lead to the idea that attention enhances the processing of task-relevant objects (“targets”) while simultaneously suppressing task-irrelevant background information (“distractors”).

In line with the idea of attention as target enhancement, attention affects not only performance but also the appearance of targets and distractors. Carrasco et al. (2004) showed that spatial attention systematically increased the perceived contrast of an attended Gabor stimulus, consistent with findings that attention increases the gain of neural responses (Maunsell, 2015). Attention also affects the appearance of several other low-level visual properties (reviewed in Carrasco & Barbot, 2019) such as gap size (Gobell & Carrasco, 2005), color saturation (Fuller & Carrasco, 2006), and motion coherence (Liu et al., 2006), mid-level properties such as shape and texture (Tian et al., 2025), and high-level properties like facial attractiveness (Störmer & Alvarez, 2016). Beyond simple enhancement, attention can also distort the perception of visual features, such as by biasing the perceived color or spatial position of a target stimulus away from distractors (Chapman et al., 2023; Suzuki & Cavanagh, 1997). Together, these findings suggest that attention biases competition towards a target by exaggerating visual features that differentiate it from irrelevant stimuli, even at the expense of accurate perception.

However, the effects of attention might not be restricted to targets and distractors, but could extend throughout the whole feature space (Chapman & Störmer, 2024). This idea follows from the framework of *representational geometry* (Kriegeskorte & Wei, 2021; Langdon et al., 2023), wherein attention-driven changes in neural information processing are best understood through their impact on the entire representation structure of the stimulus features. From this perspective, attentional selection is the process by which the representational distances among a set of stimuli are shaped to separate out the most relevant information, supporting more efficient downstream processing. These changes could take several forms. For example, when attending to a particular stimulus, such as a Gabor oriented 45° counterclockwise from vertical, changes in the gain of neurons tuned to the target orientation could distort the representational geometry (Kriegeskorte & Wei, 2021), expanding the representation around the target (orange shaded region in Figure 1). In addition, there may be compression of the representation around the distractor (purple shaded region in Figure 1). It is currently unknown whether attention modulates regions of perceptual spaces beyond target and distractor features and whether any representational expansions are offset by compression elsewhere to maintain the overall size of a representational space. Although previous work has demonstrated that attention can increase the discriminability between targets and distractors, whether and how attention alters the full representational geometry of a perceptual feature space has not been investigated.

**Figure 1.**
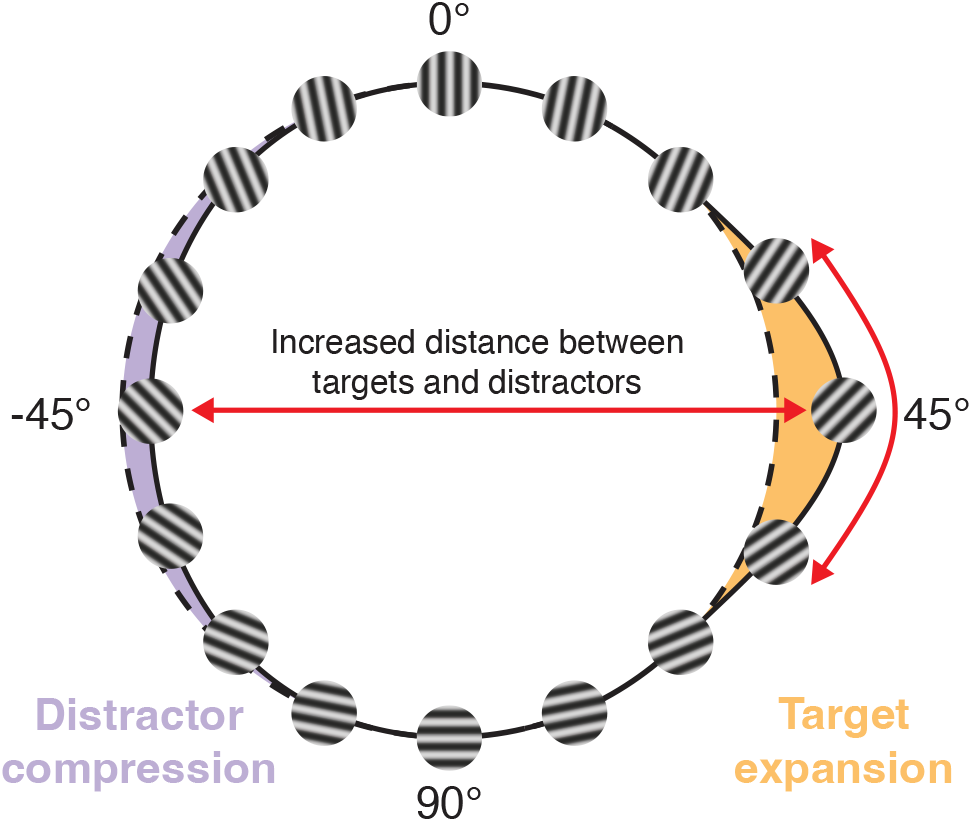
Hypothetical changes in orientation representations. When attending to a 45° stimulus, the representational geometry might expand around this part of the orientation space relative to the original representation (orange shaded region). There may also be compression around the representation of the distractor orientation (purple shaded region).

Here, we tested how attention affects the representation of orientation, a basic visual feature. Participants performed a task in which their attention was directed to particular orientations (either ± 45° relative to vertical) while we simultaneously measured the perceived similarity of orientations from around the entire feature space (Figure 2a). We modeled participants’ similarity judgments to estimate the representational geometry of orientation under these different attentional conditions, finding that attention expanded the representation in a local neighborhood around attended orientations. This expansion was accompanied by representational compression in a much broader range around the unattended orientation. The results demonstrate that the effects of attention are not isolated to changes in the processing of targets or distractors, but that attention affects the overall representational geometry of perceptual feature spaces.

**Figure 2.**
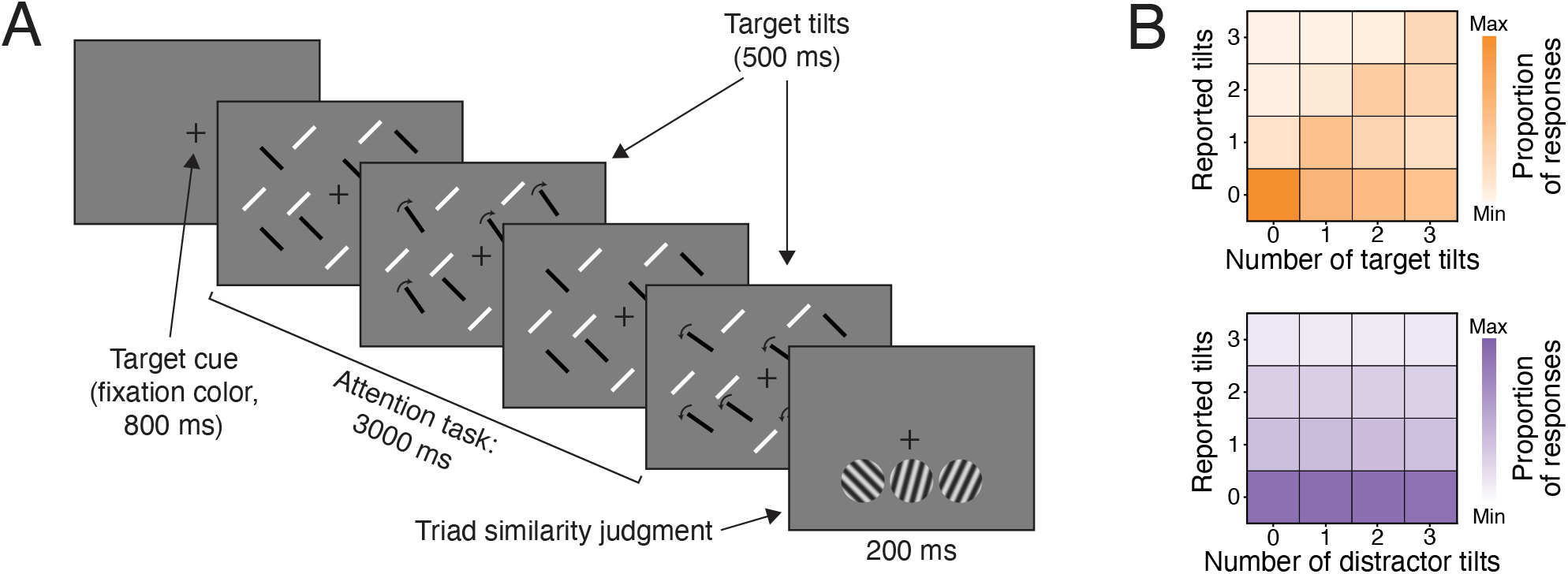
A) Example trial of the task. Participants were cued to attend to one set of oriented lines, indicated by the color of the fixation cross (black or white), and count the number of tilts in the target stimuli throughout the 3000 ms presentation period, indicated by the arrows which were not shown. At the end of each trial, participants reported which of two oriented Gabors (left or right) was most similar to the reference (center). In “no attention” trials, the attention task was not shown and participants only responded to the triad task. Stimuli are not shown to scale. B) Summary of behavioral responses in the attention task. Each cell in the grid shows the proportion of times participants reported 0, 1, 2, or 3 tilts based on the number of tilts that occurred in the target stimuli (upper) or distractor stimuli (lower). Responses tracked target tilts relatively accurately, indicated by the darker shading along the diagonal.

## Methods

### Participants

Ten participants were recruited from the Boston University community, including two of the authors (AFC and MA). All participants provided informed consent prior to participation in the study and were compensated at a rate of $15/hr (except authors). The sample consisted of 9 women and one man between 21–31 years of age (*M* = 25.2, *SD* = 3.6) with normal or corrected-to-normal vision.

### Stimuli & Apparatus

The experiment was displayed on a VIEWPixx monitor (VPixx Technologies Inc., QC, Canada) with a refresh rate of 120 Hz. All stimuli were generated and presented using PsychToolBox 3.0 (Brainard, 1997; Kleiner et al., 2007) running on MATLAB 2022a (The MathWorks, Natick, MA). Participants were seated 57 cm from the display, with head position maintained using a chinrest. The background display color was 50% grey (mean luminance = 45.56 cd/m^2^) throughout the experiment. A fixation cross was presented in the center of the display during the task.

#### Attention task

On each attention task trial, we generated an array of 100 oriented lines (50 black, 50 white, 0.75° × 5° 0.11°) that were positioned in an annulus between 1.5° and from fixation. Bars in the same color were oriented at the same angle, either 45° clockwise or counterclockwise from vertical. The starting position of the center of each bar on each trial was selected from a grid of 136 locations with 0.71° horizontal and vertical spacing. Stochastic jitter was added to the position of each bar on every frame by adding ±~0.03° to the xand y-coordinates, resulting in a random displacement of the bars across the course of the trial. The color of the fixation cross cued which set of oriented lines were to be attended, which varied across blocks.

Throughout the trial, tilt events were added to a random 80% subset of the bars in each color by adding an additional tilt offset that was titrated for each individual to achieve 70% accuracy (described below). On each trial, there were either 0, 1, 2, or 3 tilt events in each color, and each tilt event was randomly selected as clockwise or counterclockwise relative to the base orientation. Each tilt lasted 500 ms and events could not occur within 200 ms of the beginning or end of each trial, or within 200 ms of a previous event in the same color (though tilts in different colors could overlap in time).

#### Triad similarity task

Triad task stimuli were oriented Gabors, 2° in diameter within a Gaussian envelope (0.2° SD) at a spatial frequency of 4 cycles/° and a contrast of 64%.

Three Gabors were shown, each with a 1° vertical offset below fixation. The middle Gabor was centered horizontally, while the other two were offset by 2° to the left or right. Each Gabor was oriented at a different angle, sampled without replacement from the full range between 0-175° at 5° spacing, generating 36 unique orientations.

### Procedure

Participants completed the experiment across several sessions. In the first session, they were trained on the attention and triad tasks separately, before completing a shorter run of the full task. In the next 4-5 sessions, they completed longer runs of the full task. Participants also completed separate “triad only” sessions, in which they performed the triad task alone, which we used to assess the representational structure of orientation independent of attention. Seven participants completed the triad only sessions as their final sessions, while three participants completed them as their first sessions.

The attention task was designed to engage feature-based attention towards a particular orientation. Participants were cued to attend to bars in a particular color (black or white) that were oriented 45° clockwise (“attend 45°”) or counterclockwise (“attend −45°”) from vertical. The fixation cross was presented in the target color for 800 ms before each trial.

During each trial (3000 ms in duration), bars in each color tilted clockwise or counterclockwise 0–3 times by an additional offset. Participants were instructed to count the number of tilts in the attended color, to report at the end of the trial (using the numbers 1–3, or the ‘~’ key for 0, with their left hand).

The triad similarity task was designed to measure the perceptual similarity between orientations. At the offset of the attention task, the triad similarity task stimuli, an array of three Gabors, were presented in a row below fixation for 200 ms. Participants were instructed to report which of two Gabors (‘9’ for left, ‘0’ for right, with their right hand) was most similar in orientation to the center Gabor. To prioritize triad judgments based on perceptual similarity, participants reported them before their tilt response from the attention task. The full timeline of each trial was: 1) attention task stimulus, 2) triad stimulus, 3) triad response, 4) attention task response. The responses to each task were unspeeded, and response accuracy was emphasized.

At the beginning of each session, participants completed a thresholding task to determine the magnitude of the tilt event offset. The thresholding task consisted of 64 trials similar to the main attention task, except trials were 2000 ms in duration and only a single target and distractor tilt event was presented on each trial. Participants were instructed to report the direction of the tilt, relative to the orientation of the target bars (‘1’ = counterclockwise, ‘2’ = clockwise). The initial tilt offset was set to 20° and was adjusted using a 3-down, 1up staircase procedure (i.e., after three correct responses, the tilt offset was reduced). The possible tilt offsets were 20°, 16°, 12°, 8°, 6°, 4°, 3°, 2°, and 1°. We defined the thresholded tilt value as the mean tilt across the last 5 reversals during the staircasing procedure. Convergence of the procedure was determined by plotting the tilt offsets across trials and comparing this to the threshold estimate. Thresholds ranged from 1.6°-6.4° across sessions, with a mean of 3.05° (*SD* = 0.98).

The attention task was divided into blocks of trials in which the target orientation and color was constant. Participants completed 384 or 480 trials per session, separated into eight blocks (48 or 60 trials per block) such that each combination of target orientation and color was repeated twice. The number of target and distractor tilt events was drawn randomly on each trial, such that 0, 1, 2, or 3 tilts was equally likely, and each event was randomly determined to be a clockwise or counterclockwise tilt. The orientation of each triad task stimulus was sampled without replacement from the set of 36 orientations.

For triad only sessions, no thresholding procedure was required. In each session, participants completed 640 trials of the triad task, sorted randomly into 8 blocks of 80 (except one participant, who in one session completed 480 trials in 8 blocks of 60). This resulted in a roughly equal number of triad responses across sessions for each condition (attend 45°, attend − 45°, triad only): we obtained 1152-1776 (*M* = 1339.2, *SD* = 188.6) trials for each condition in the attention task, and 1120-2560 (*M* = 1584, *SD* = 556.6) trials in the triad only task.

### Analysis

#### Attention task analysis

For each participant and for each target orientation, we first summarized behavioral performance by calculating the number of times each response was made (0, 1, 2, or 3 tilts) as a function of the presented number of target tilts. Accuracy in the attention task was calculated as the proportion of times participants reported the exact number of target tilts. However, because we thresholded participants based on tilt discrimination, such that each target tilt was detected with approximately 70-80% accuracy, reported tilts were expected to be lower on average than the number of presented target tilts. To better quantify behavior in the attention task, we therefore also computed Spearman’s correlations between the reported and presented tilts for each participant. As a comparison, and to confirm that participants attended selectively to the cued target orientation, we also calculated correlations between the reported target tilts and the number or presented distractor tilts.

#### Representational dissimilarity matrices

To assess the perceptual similarity of orientations in the triad similarity judgment task, we first estimated the *representational dissimilarity matrices* (RDMs) based on behavioral responses. On each trial, participants report which of two stimulus orientations they perceive as most similar to the central reference orientation. For each participant and each attention condition, we constructed 36 × 36 RDMs such that the value in each cell corresponded to the proportion of trials in which a particular combination of orientations was chosen as most similar when they were presented together, regardless of the orientation of the third stimulus. The grand average RDM is presented in Figure 3a.

**Figure 3.**
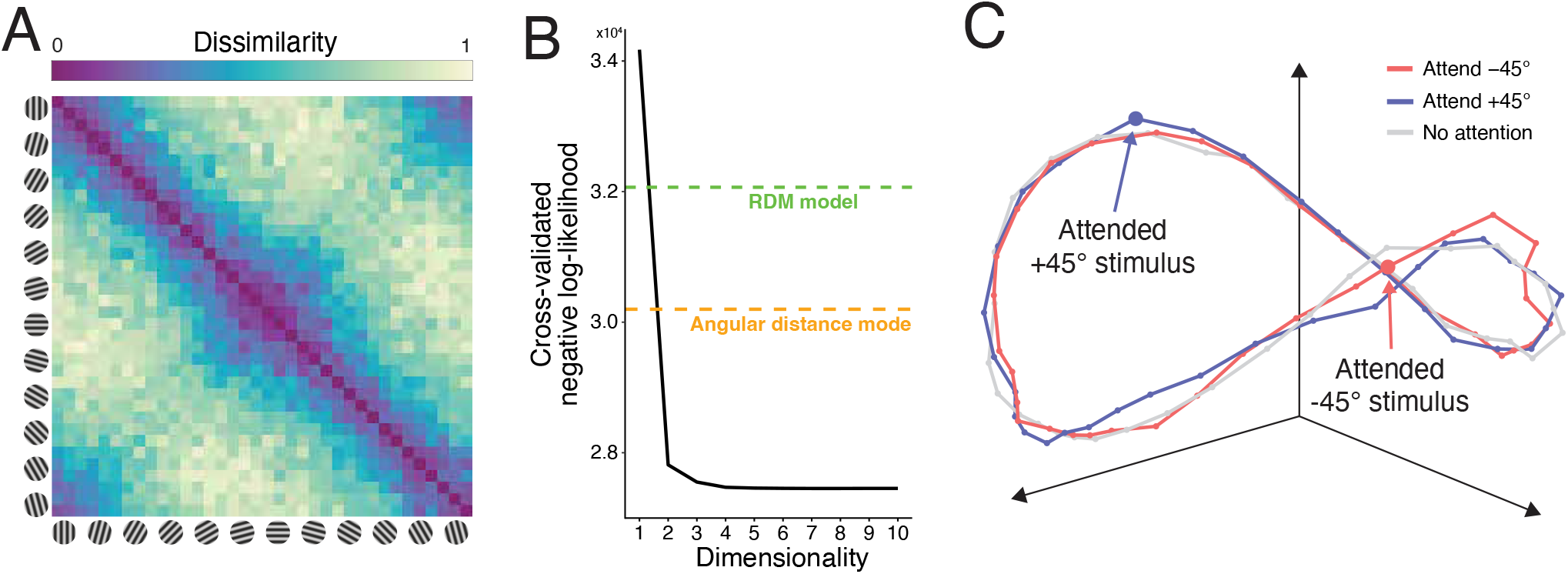
A) Grand average representational dissimilarity matrix across all trials in the similarity judgment task. Each cell in the matrix indicates the dissimilarity between a given pair of oriented Gabors, which we calculated as the complement of the proportion of times that pair was chosen as most similar when presented together. B) Cross-validated negative log-likelihood of the best fitting behavioral model at each dimensionality. Model fits improved up to around 4 dimensions, at which point the negative log-likelihood remained nearly constant. Horizontal lines indicate the fit of two additional models: the “angular distance model” in which the model predicted behavior only on the basis of the difference in orientations between triad stimuli, and; the “RDM model” in which we fit the dissimilarities directly, ignoring geometric structure. C) Group-average 3-dimensional solution for each attention condition. The representational structure of the similarity judgments approximately followed a saddle shape in the 3-D space. Coordinates corresponding to the two target orientations (± 45°) showed deviations as a function of the attention conditions.

#### Representational geometry of similarity judgments

To estimate the representational geometry of orientation from participants’ similarity judgments, we implemented a variant of multidimensional scaling developed by Waraich and Victor (2024). The method seeks to place the stimuli in an *m*-dimensional coordinate space, such that the distance between stimuli in this space is predictive of the perceived (dis)similarity between them. We modeled decisions in the triad similarity judgment task as comparisons of the distance between the reference, *s*_*re f*_, and the two choice stimuli, *s*_*left*_ and *s*_*right*_. Assuming that uncertainty in decisions is affected by additive Gaussian noise, the probability of choosing the left stimulus as being most similar to the reference is modeled by the function:

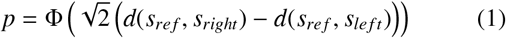

where Φ is the standard normal cumulative distribution function and *d*(·,·) is the Euclidean distance between two stimuli (though, in general, other metrics could be used).

The log-likelihood of the observed similarity judgments across trials is:

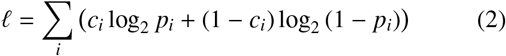

where *c*_*i*_ is an indexing variable for each trial *i*, such that *c*_*i*_ = 1 when the left stimulus was chosen, and *c*_*i*_ = 0 when the right stimulus was chosen, and *p*_*i*_ is the expected probability of choosing the left stimulus as being most similar to the reference given the stimuli on trial *i*, as calculated in Equation 1.

#### Model fitting

The model parameters are the *m*dimensional stimulus coordinates, which determine the pairwise distances between stimuli, and therefore the choice probabilities, *p*. The coordinates are adjusted across iterations to minimize the negative log-likelihood of the triad similarity judgments. We fitted this model to data from all participants with *f minunc* in MATLAB (2024a), using the quasiNewton method and L-BFGS approximation of the Hessian (memory = 20). To aid in optimization, we computed the analytical gradients of the negative log-likelihood function (see Appendix A).

We fit a series of models aimed to assess the dimensionality of the representational geometry of orientations, reflected in the behavioral similarity judgments, as well as the effects of attention. The coordinates were represented in a 4dimensional tensor ***X***_*i jkl*_, with each cell representing the coordinate of stimulus *i* along dimension *j* in attention condition *k* for participant *l*. For all models, we added two separate penalties to the negative log-likelihood to constrain fits.

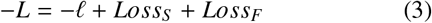

The group ridge penalty, *Loss*_*S*_, was added based on the deviations of coordinates between participants:

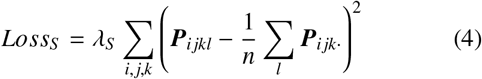

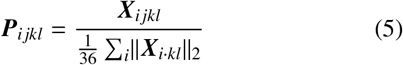

where *λ*_*S*_ is the ridge parameter that scales the amount of penalization in the model. This penalization has two primary effects on model fits: 1) individual participant coordinate spaces are shrunk towards the group mean; 2) coordinate spaces from different participants are aligned, allowing for direct comparison. ***P*** is a rescaled coordinate space for each participant based on the mean distance of the stimuli from the origin, so that participants with coordinate spaces that are larger or smaller than the mean are not excessively penalized.

We also applied a ridge fusion penalty, *Loss*_*F*_, which penalized large deviations between stimuli that represent similar orientations (Goeman, 2008):

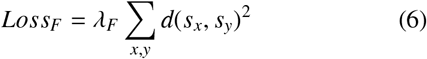

given that *s*_*x*_ and *s*_*y*_ are neighboring stimuli (i.e., 5° apart). This penalty imposes smoothness of the resulting coordinates.

The best fitting model for a given dimensionality was selected using 10-fold cross-validation. We first divided the data from the triad task into 10 sets, balanced so that each set contained nearly equal numbers from a given participant and attention condition (though within each set, trial numbers did differ between participants). To determine the initial point for the optimizer, we first constructed the group average RDM of the training data (described in **Representational dissimilarity matrices**) and performed non-metric multidimensional scaling using *mdscale*. This solution was then used to fit two models collapsed across attention conditions, one with shared coordinates across all participants and one that varied for individual participants. We then averaged the coordinates of these fits, to get an initial point that was halfway between the group coordinate space and individual spaces.

To fit the main models, for each combination of *λ*_*S*_ and *λ*_*F*_ values, we used 9 folds as model training data to extract the best fitting coordinates and then tested the model on the last held-out fold to obtain the cross-validated negative log-likelihood. We iterated this process such that each set was used as test data once per pair of ridge parameters, and summed the negative log-likelihoods across all folds. Given that some participants completed more trials than others, we weighted the negative log-likelihood of each fit such that each participant’s data contributed equally to model fits.

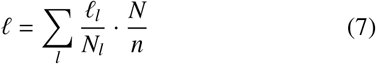

where *ℓ*_*l*_ is the log-likelihood of the model for participant *l, N*_*l*_ is the number of trials completed by that participant, and *N* is the total number of trials in the dataset.

Across different model fits, *λ*_*S*_ was set to zero or one of 20 values log-spaced between 10^−1.5^ to 10^3^, while *λ*_*F*_ was set to zero or one of 20 values log-spaced between 10^−1.5^ to 10^2.5^. For each level of dimensionality, we selected the best fitting model with the values of *λ*_*S*_ and *λ*_*F*_ that minimized the cross-validated negative log-likelihood.

We fit all models twice, first assuming no differences as a function of attention condition (i.e., *k* = 1, requiring 36 × *m* × 10 parameters, for stimuli, dimensions, and participants, respectively) resulting in coordinate spaces that reflect the average for each participant across the full dataset from the triad task. Second, to assess how attention modulated the coordinate spaces, we fit the model with different coordinates for each attention condition (i.e., *k* = 3, requiring 36 × *m* × 3 × 10 parameters) using an otherwise identical modeling procedure.

#### Alternative models

We tested our main geometric model against two alternatives. Both models were fit using 10-fold cross-validation and were optimized to minimize the same negative log-likelihood as the main model. The only change was in how we defined the distance between pairs of orientations.

The first alternative model was the “angular distance model”, in which the distances between pairs of orientations was defined as the arc length along the circumference of a circle spanning the orientation space. This model reflects a lower bound on geometric complexity.

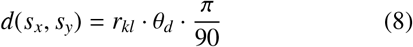

where θ_*d*_ is the difference in orientations between *s*_*x*_ and *s*_*y*_ (in degrees), and *r*_*kl*_ is the radius of the circle for participant *l* and attention condition *k*. We fit this model with the radii as free parameters, requiring 30 parameters (3 per participant) when fitting each attention condition, or 10 parameters when collapsed across conditions. We initialized the model parameters as a vector of ones.

The second alternative model was the “RDM model”. Here, we eschewed the geometric structure of the main model, allowing each cell of the model RDM to vary as a free parameter. This model provides an upper bound on complexity and is more data-driven. We assumed a symmetric RDM with zeros along the main diagonal, meaning there were 36 × 35/2 = 630 parameters per individual RDM, for a total of 18,900 parameters for fitting separate attention conditions, or 6,300 when collapsed across condition. With this large number of free parameters, we attempted to reduce the effects of overfitting by applying a ridge penalty to the sum of the squared model parameters. We set the model *λ* to zero or one of 20 values log-spaced between 10^−3^ and 10^3^, selecting the model that minimized the cross-validated negative log-likelihood. The initial model parameters were set by constructing the RDM based on the training data, as in the original main model, then taking the values in the lower triangle.

#### Assessing the effects of attention on representational geometry

After extracting the representational geometry underlying perceptual similarity judgments in the triad task, we computed the length of the representational space in a local neighborhood around each orientation, which we refer to as the *local length*. We selected stimuli in a window spanning ± 20° around each orientation, then computed the sum of the Euclidean distance between neighboring stimuli within this window. This approach provides an estimate of the distance along the curve in the *m*-dimensional representational space near each orientation, following previous modeling work (Ringach, 2010, 2019). We used a window of ±20° since this resulted in four non-overlapping bins centered at the cardinal (0° and 90°) and oblique (±45°) orientations, with each stimulus orientation falling into one of these bins.

We normalized the length in each bin by dividing it by the total length spanned by all orientations, separately for each participant and attention condition, such that local lengths reflect a proportional measure of the representational space near a given orientation. This allows us to better compare changes in the perceptual geometry around the attended and unattended orientations, and during the triad only sessions where attention was not directed to a specific part of the feature space, independent of variations in the overall size of the representational geometries between participants and conditions. For key results, we also performed the same analyses using the original (non-normalized) lengths.

For statistical comparisons, we selected the local lengths at ±45° when each orientation was the target, distractor, or when no attention was directed towards it. These lengths were then submitted to a two-way repeated measures ANOVA. In later analyses, we selected the local lengths at the cardinal and oblique orientations, and conducted a twoway repeated measures ANOVA to assess the interaction between these orientations and the attention conditions. In all cases, significant effects were followed up by paired-samples *t*-tests comparing lengths at a given orientation between the attention conditions directly.

We also analyzed how local lengths changed in each attention condition as a function of the window size used to calculate lengths. Increasing this window naturally increases the overall length, so we divided all lengths by the size of the window, resulting in a measure of the mean distance between neighboring stimuli across the window. We then directly assessed the difference in lengths between target and distractor orientations, for different combinations of each window size. We limited the window size combinations to avoid cases in which any stimulus fell into both windows (i.e., target window + distractor window ≤ 180°). To assess significant differences between lengths across different combinations of window sizes, we conducted permutation tests to determine a significance threshold. We refit the 4-D geometric model while randomly shuffling the attention condition labels across trials for each participant 10,000 times. We used the best fitting ridge parameters determined for the original model, and extracted new model coordinates with each shuffle of the permuted data. For each permutation, we calculated local lengths for different window sizes as above, and determined the maximum absolute difference between (shuffled) target and distractor lengths. We identified significant window combinations in the original data if the difference between lengths was greater than 95

## Results

### Attention to target orientations improved tilt detection

To engage feature-based attention, participants were cued to attend to a set of target lines oriented 45° clockwise or counterclockwise from the cardinal axes. During each trial, both target and distractor lines could tilt slightly away from their main angle 0–3 times, and participants were instructed to count the number of tilts that occurred in the target lines, while ignoring tilts in the distractor lines.

We first confirmed that participants were able to detect and report target tilts, finding that average accuracy was 52.5% (*SD* = 13.4), significantly exceeding 4-AFC chance performance of 25%, *t*(9) = 6.50, *p* <.001. Participants tended to underestimate the number of target tilts, with a mean response of 1.23, when the empirical average was 1.50. This underestimation was reflected in the distribution of participants’ responses, which were skewed towards reporting zero, one, or two tilts (26.1%, 34.6%, and 29.2% of trials, respectively), rather than three tilts (10.2% of trials). Such underestimation was expected, as the size of the tilt was thresholded for each participant to achieve 79

We next assessed the effectiveness of feature-based attention in this task by examining how responses varied as a function of the number of target and distractor tilts. Participants’ reports generally tracked the number of target tilts, as shown by positive correlations between the presented and reported number of target tilts, *r*_*mean*_ =.646 (Figure 2b, upper). In contrast, correlations between responses and the number of distractor tilts were near zero across participants, *r*_*mean*_ =.009 (Figure 2b, lower). This pattern of behavior confirms that participants attended to the target orientation, as instructed, as responses were systematically related to tilts in the target-oriented lines only.

### Extracting the representational structure of orientation

To measure the full representational geometry of orientation and how this structure changes with attention, it is necessary to measure the appearance of all orientations, and not just the target and distractor orientations presented during the attention task. To achieve this complete measurement of perceptual orientation space, participants performed a triad similarity judgment task interleaved with the attention task. On each trial, an array of three oriented Gabors was presented for 200 ms, and participants were instructed to select which of two stimuli (the leftmost or rightmost) was more similar in orientation to the third, reference stimulus (in the center; Figure 2a). The orientation of each stimulus was randomly and independently sampled from 0° to 180° with 5° spacing. We collected data from this task under three attention conditions: either the triad task performed in isolation (“no attention”), or when attention was directed to 45° or −45°. We hypothesized that any changes to perceptual geometry during the attention task on each trial would affect the appearance of the triad probe stimuli presented immediately afterward, allowing us to measure the representational changes induced by attention.

We first sought to characterize the perceptual geometry of orientation regardless of attention by combining data across the three attention conditions. To quantify the representational structure of orientation, we computed representational dissimilarity matrices (Figure 3a), which summarized the proportion of times participants selected a particular pair of orientations in each triad as being most dissimilar. The group average RDM demonstrates the expected low dissimilarity near the primary diagonal where the angular difference in orientations is smaller (e.g., only 5° or 10° apart), and high dissimilarity off the main diagonal peaking at orthogonal angles. We also observed variability in the RDMs at the level of individual participants (see Figure S1), suggestive of individual differences in similarity judgments.

To estimate the representational structure of orientation, we fit a geometric model to similarity judgments in the triad task. The model seeks to find a set of coordinates that predicts behavior, such that the distance between stimuli in the *m*-dimensional representational space reflects the perceived similarity between them. We measured the model’s performance using the cross-validated negative log-likelihood of model fits as we varied the dimensionality of the representational space. We first assessed the dimensionality of the perceptual representations using a single set of coordinates across attention conditions, finding that while each additional dimension continued to decrease the negative loglikelihood up to the maximum dimensionality we examined (*m* = 10), there was little improvement beyond 4 dimensions (Figure 3b). For higher-dimensional fits, the additional dimensions had very low variance and in some cases were correlated, suggesting they contained little explanatory power compared to the first four dimensions.

We compared these geometric models to two additional models. In the angular distance model, behavioral predictions were based on the differences in orientation between the two triad stimuli and the reference (equivalently, the arc length along a circle between two orientations). This model was only an improvement over the 1-D geometric model (orange dashed line in Figure 3b), indicating that similarity judgments were based on more than just the angular deviations between Gabors. In the RDM model, we fitted the dissimilarity values for all pairs of orientations directly, allowing for a characterization of behavior that was not constrained to a particular geometry. However, this model also failed to provide a better fit to the data (green dashed line in Figure 3b), performing even worse than the angular distance model. While the fitted RDMs showed some correspondence to those measured directly from behavior, they appeared to overfit to the training data, resulting in worse out-of-sample predictions. Thus, the constraints on distances imposed by the structure of the geometric models resulted in better generalization, suggesting these models captured meaningful representations of the perceived similarity between orientations.

The individual dimensions of the 4-D solution are shown in Figure S2a. The first two primary dimensions captured the circular structure of the stimulus space, approximated by sin and cos functions. The third and fourth dimensions were approximated by the second harmonics of the first two dimensions, together forming a “double loop” structure where orientations differing by 90° are represented similarly in these dimensions (Figure S2b). Another way of interpreting the higher dimensions is that they deform the circular space of orientations into a saddle-like structure. One consequence of this structure is that the perceived similarity between orientations falls off exponentially as a function of distance in the physical feature space, consistent with proposed laws of psychological similarity (Shepard, 1987; Sims, 2018). Finally, we found that the stimulus coordinates fell near a hypersphere: the distance of each point from the origin was close to uniform across the feature space. Orientations near horizontal and vertical were slightly farther from the origin than obliques, consistent with a small cardinal bias (Figure S3). Thus the perceptual space we recovered for orientation was a nonlinear but systematic transformation of the physical feature space into a higher-dimensional representation that accounts for several aspects of orientation perception.

### Attention expands representational space around target orientations

To assess how attention to different orientations modulated representational geometry, we refit the models, allowing the coordinates to vary separately for each of the three attention conditions (attend +45°, attend − 45°, no attention). We assessed the dimensionality of the representations using cross-validated log-likelihood, finding again that 3-4 dimensions provided good fits to the behavioral data (Figure S4). Compared to the models fit to the aggregated data, there was a smaller improvement for 4relative to 3-dimensional fits, possibly due to the smaller number of trials when fitting each condition separately. Our analyses focused on the 4dimensional solution, though for visualization we plotted the 3-dimensional solutions for each attention condition overlaid, observing that while the overall saddle-shaped structure was similar in all conditions, the representational geometry for different attention conditions diverged near the target orientations, particularly around ±45° (Figure 3c).

To quantify the changes in representational distance induced by attention, we computed a “local length” metric around key stimulus orientations. We divided the space into four quadrants, with 40° (±20°) windows around the target orientations of ±45° and the task-irrelevant control orientations of 0° and 90°. We then calculated the length along the representational space within the window (e.g., 25° to 65°, centered on 45°) by summing the distances spanned by neighboring points. To account for differences in the overall size of the representational space between participants or conditions, we normalized the local lengths, expressing them as a proportion of the total length around the orientation space (see Methods).

We first focused on the target-centered windows, assessing the local length when each orientation was the target, when it was the distractor, and on triad-only trials, when attention was not directed to a particular orientation. We found that attention modulated the local lengths within these windows, as shown by a main effect of attention, *F*(2, 18) = 7.03, *p* =.006, with the longest lengths around a stimulus orientation when it was the attended target orientation, the shortest lengths when it was the unattended distractor orientation, and intermediate lengths under no attention (dashed vertical lines in Figure 4a). There was no effect of stimulus orientation, *F*(2, 18) = 0.79, *p* =.396, and no interaction between attention and stimulus orientation, *F*(2, 18) = 0.60, *p* =.562, suggesting that the effects of attention were similar at +45° and − 45°. Pairwise comparisons between attention conditions revealed a significant increase in the lengths around targets relative to distractors, *t*(9) = 4.16, *p* =.002, with a marginal increase around targets compared to no attention, *t*(9) = 2.22, *p* =.053. There was no difference in lengths for distractors relative to no attention, *t*(9) = 1.10, *p* =.302. These results indicate that attention expands the representational space in a local window around target stimuli.

**Figure 4.**
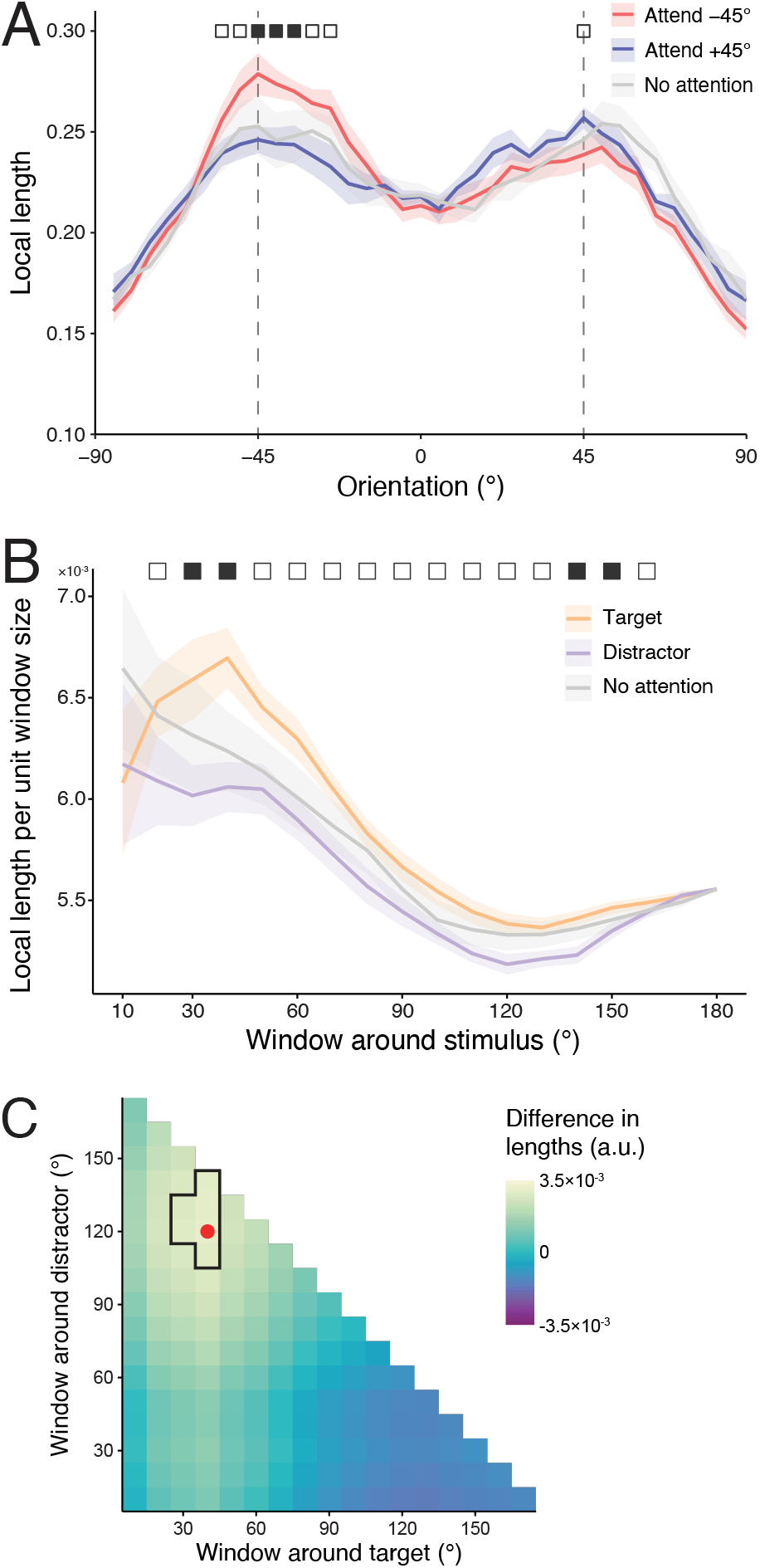
A) Local lengths of perceptual representations of orientations in each attention condition. Shaded regions indicate within-subjects SE. Significant differences in lengths between “Attend − 45°” and “Attend +45°” are indicated by filled squares (Bonferroni corrected across 36 orientations, p <.05) or unfilled squares (uncorrected p <.05). B) Local length around ± 45° orientations scaled by the size of the window around this orientation and whether it represents the target or distractor orientation, or when no orientation was attended. C) Window sizes maximizing relative expansion around targets vs. compression around distractors. The outlined region shows combinations of targetand distractorcentered windows with significant differences in lengths between conditions (permutation test p <.05), with the red dot indicating the windows with the largest observed difference.

Next, to investigate how attention affected the representation of orientations throughout the stimulus space, we compared local lengths in windows centered on each of the cardinal (0° and 90°) and oblique orientations (+45° and −45°) as a function of the attention condition. We found that attention had different effects across the representational space, indicated by a significant interaction between attention condition and stimulus orientation, *F*(6, 54) = 2.64, *p* =.026. Assessing each quadrant window separately, we found a significant increase in the lengths around − 45° when attention was directed to that orientation, *F*(2, 18) = 3.78, *p* =.043, particularly when compared to attending to +45°, *t*(9) = 4.89, *p* <.001. At +45°, there was no main effect of attention, *F*(2, 18) = 2.68, *p* =.096, but attending to +45° vs − 45° resulted in significantly greater lengths, *t*(9) = 2.54, *p* =.032. When we compared lengths around 0° or 90°, there were no significant differences due to attention, *p*^′^s >.3, demonstrating that the effect of attention was specific to the target orientations at ±45°.

Lastly, we conducted a sliding window analysis, computing the local length in windows (width = 40°) centered at each stimulus orientation (Figure 4a). Inspection of the resulting local length plot showed that the effects of attention were largest around −45°, with a smaller, but still visible effect observed at +45°. A similar pattern was observed using the original (non-normalized) local lengths (Figure S5a). Thus, attention modulated the representational structure of orientation, specifically by increasing the local lengths around attended orientations, hence expanding the relative representational space near that point.

### Attention compresses representations broadly around distractor orientations

If attention expands the representation around targets, does this imply that attention can increase the overall size of representational spaces? To address this question, we computed the total (non-normalized) length spanned by the full orientation space in each attention condition. These total lengths did not significantly differ, *F*(2, 18) = 0.14, *p* = 0.87, suggesting that attention did not expand the available space for representing orientation information. In fact, the total length was the greatest numerically under no attention. This finding suggests there were subtle *decreases* in the lengths around unattended orientations that occurred alongside the increases at the attended orientation.

Examination of Figure 4a suggested that representational compression around distractors may be present but both smaller in magnitude and less focal than representational expansion around targets. To investigate this possibility, we varied the size of the window around the target and distractor orientations and calculated local lengths for each window size in each attention condition (Figure 4b). There were significant differences between target and distractor lengths for a wide range of window sizes, though the effects peaked around 30–40°, which was mirrored at 140–150° due to symmetry across the space when collapsing across the two target orientations. A similar pattern was observed for the original lengths (Figure S5b). Numerically, we found both expansion (longer lengths around target orientations vs. no attention) and compression (shorter lengths around distractor orientations vs. no attention) across a wide range of window sizes, for both the normalized (Figure S6a) and original lengths (Figure S6b). Whereas the expansion around the target peaked for narrow windows of 30–50°, compression was most reliable for wider windows of 110–130°. This result implies a broad compression of the representational space that encompasses a range of stimulus orientations spanning the distractor as well as other non-target orientations.

To better characterize the potentially different breadths of expansion and compression in the perceptual space, we recalculated the difference in lengths around target and distractor orientations while allowing the size of both windows to vary (without overlap). We observed significant differences in lengths for a number of window combinations, with the largest effect of attention occurring for a target window of 40° and a distractor window of 120° (Figure 4c, see also Figure S5c). Thus, our findings suggest that attention induces representational expansion in a narrow range around the target and compression in a broad range around the distractor.

## Discussion

In this study, we examined how attention affects representational geometries, using perceived similarity to model the perceptual space of orientation (Roads & Love, 2024; Zaidi et al., 2013). Participants performed a task in which they directed attention towards one of two sets of oriented lines (±45° from vertical) to detect small tilt offsets in the target stimuli, while at the end of each trial they performed a perceptual similarity judgment among three oriented Gabors, independently sampled from the 180° stimulus space. We fit a model to the similarity judgments to extract the representational geometry of orientation under different attention conditions, finding that orientation was best described by a 4-dimensional model. From the fitted representational structure, we observed a reliable expansion in a region spanning the attended orientation compared to when it was unattended. We estimated the breadth of attentional expansion for orientation to be approximately 40°, as this window size maximized the attention-induced change in representational distances around the target. In contrast to this relatively narrow expansion, we found a broad compression of the representational space around distractors that peaked with a window of 120°. Thus, our findings show that the effects of attention are not confined to the specific target and distractor features in a particular task, but that attention reshapes the entire representational geometry of a given feature space (as hypothesized by Chapman & Störmer, 2024).

Our behavioral models showed that the perceptual representation of orientation is best captured in 4 dimensions, higher than the physical 2-D geometry of the circular feature space. Indeed, a simple angular distance model performed substantially worse than the 4-D geometric model. Interestingly, this is consistent with findings that the perceived similarity between the orientation of abstract shapes is also well described by four dimensions (Shepard & Farrell, 1985). In particular, while two dimensions observed by Shepard and Farrell (1985) captured the basic circular structure of orientation, the other two dimensions had a “double-loop” structure where orientations differing by half a cycle were represented more similarly. Recent neuroimaging work suggests that the sensory representation of orientation has a more circular structure in early visual cortex (V1–3) that is transformed in higher-order visual regions (LO) into a structure that may be consistent with a double-loop (Chunharas et al., 2025). The 4-D similarity structure we observed thus suggests that the perceptual representation of orientation, even for simple Gabor stimuli with no complex shape or object structure, may be an outcome of the successive stages of processing along the visual hierarchy.

Notably, our data support predictions of previous theoretical work on the representational geometry of basic, circular feature dimensions (Ringach, 2010). Ringach (2010) determined several properties of the optimal representation of such features that would hold under the conditions that 1) the population undergoes normalization and 2) the neuronal tuning functions are translation invariant (i.e., they symmetrically and uniformly tile the stimulus space). One theoretical result was that the representational manifold produced by this model lay on a hypersphere, due to normalization. Our findings showed not only that the orientation representation as a whole lay near a hypersphere, but that attention did not change the overall size of the representational space, with expansion and compression cancelling out, consistent with normalization. A second important theoretical result was that, in such optimal neural representations, successive pairs of dimensions produce loops with increasing harmonics. We found the predicted single- and double-loops in our main 4-D model, and further our higher dimensional models show evidence for a triple-loop at dimensions 5 and 6, which to our knowledge has not previously been reported (Figure S7). Higher-order loops did not appear for even higher dimensions, suggesting an upper limit on the complexity of the representation of orientation (Ringach, 2010), although our ability to determine these dimensions from our data may have been limited. However, we find that these loops were not symmetric in our data, with separation still observed near cardinal orientations in the double-loop. This asymmetry may be because orientation tuning is not fully *translation invariant* (as the model of Ringach, 2010, requires), with potential differences in tuning around cardinal compared to oblique orientations (Girshick et al., 2011; Wolff & Rademaker, 2025). Further work will be needed to determine the extent to which the higher dimensions of these representations contribute to orientation perception.

Critically, we found that attention affected the overall structure of this representational space. Although data from previous studies has not allowed measurement of whether and how attention warps the full representational space, classic work has demonstrated how different dimensions of stimulus representations can be flexibly weighted more or less strongly in perceptual decisions. This idea hearkens back to Attneave (1950), who showed that participants’ judgments of the similarity between simple geometric shapes were consistent with a city-block metric, such that decisions reflected a weighted sum along individual stimulus dimensions. Subsequently, Shepard (1964) argued that stimuli with analyzable (i.e., separable or independent) dimensions are compared primarily along one dimension or another, resulting in distances that do not conform to a Euclidean metric. In contrast, unitary stimulus spaces are perceived more holistically and, Shepard proposes, reflected in distances consistent with a Euclidean metric. The notion of the independence of stimulus dimensions is embedded in multidimensional scaling methods, such as INDSCAL (Carroll & Chang, 1970), where individual differences are accounted for by applying separate weights to each dimension in the model. Such models can only account for linear deviations between individuals or conditions, and thus are unable to capture the differences in the orientation representations we observed, which were not isolated along specific dimensions but rather were captured by a non-linear warping across multiple dimensions simultaneously. Thus, our data support the idea that orientation is a unitary feature, requiring a methodological approach different from multidimensional scaling to measure changes in its representational structure. This conclusion is further supported by recent work showing that Euclidean geometries can adequately capture the perceptual and semantic similarity of visual stimuli (Waraich & Victor, 2024), in contrast with the hyperbolic geometry found in the olfactory system (Zhou et al., 2018).

Our findings support the hypothesis that attention reshapes perceptual representations in a way that supports the processing of task-relevant information (Chapman & Störmer, 2024). In our task, participants were cued to attend to sets of lines oriented 45° clockwise or counterclockwise from vertical, and to detect small tilts in these lines throughout the trial. Attention-induced changes in representational geometry can enhance performance by increasing the perceived distances between orientations nearby the target. That is, tilts in the target stimuli (e.g., from − 45° to −47°) are made more discriminable because attention increases the representational distance between these orientations, effectively exaggerating the tilts. This interpretation aligns with previous findings, whereby attention affected the perception of stimulus properties, such as contrast (Carrasco et al., 2004) or gap size (Gobell & Carrasco, 2005), which can allow for enhanced discriminability of attended relative to unattended stimuli. We observed clear differences in perceptual representations when attention was directed to specific orientations (attend 45° vs. attend −45°), allowing us to measure expansion vs. compression relative to the size of the whole space, but less reliable differences when comparing between attention and no attention conditions, particularly when using non-normalized lengths, making claims of absolute expansion or compression of these representations less certain. Comparisons of attention vs. no attention may have been less reliable both because these differences are inherently smaller than target vs. distractor differences and because the no attention data was collected in separate experimental sessions. Future studies should aim to better estimate changes in absolute representational geometry relative to a neutral baseline in addition to relative changes across the space.

Other work suggests that attention may not just exaggerate task-relevant features, but can also distort perception. Attention can bias the perception of spatial position (Suzuki & Cavanagh, 1997) and color hue (Chapman et al., 2023), where the magnitude of the hue bias depends on the similarity between target and distractor colors. This finding suggests that attention can alter perceptual representations in different ways depending on context: when target and distractors are dissimilar (such as the orthogonal orientations in our attention task), attention acts to increase the representational distances around the target feature; in contrast, when targets and distractors are more similar, attention may act to distort the representation of target features *away* from distractors, resulting in biased perception of the target itself. Both mechanisms are consistent with the broader idea that attention reshapes representations to allow for more efficient transfer of perceptual information that is relevant for behavior (Ruff & Cohen, 2019; Rust & Cohen, 2022). Thus, while the expansion and compression we observed are ways in which attention can reshape representational spaces, other mechanisms are plausible, and task context likely has a large effect on what information is magnified by attention (Kay et al., 2023). Representational geometry has often been studied using neural measures, allowing for researchers to examine how representations change across brain regions involved in perception and memory (Chunharas et al., 2025; Xu, 2023), or how representations are affected by visual masking (Ringach, 2019), for example. In comparison, behavioral measures of similarity reflect the outcome of both perceptual and decision-making processes. Given that the dimensions underlying object representations vary along the visual hierarchy (Contier et al., 2024), we expect that attention might affect the representational geometry in different ways at different stages of processing. Likewise, attention may impact the temporal evolution of feature representations during visual processing. In our task, we assessed perceptual similarity only at the end of each trial, after attention had been directed to the target orientation for several seconds. Neural measures with high temporal resolution can instead be used to track representations during early visual processing, as demonstrated by recent work showing that spatial position (Foster et al., 2021), color (Rosenthal et al., 2021), and even semantic object properties (Teichmann et al., 2025) can be recovered from multivariate EEG/MEG activity. Combining perceptual similarity judgments with neural measures can therefore provide key insights into the way attention shapes stimulus representations during visual processing, as well as providing a powerful way to determine which neural signals map to different aspects of perception and cognition. More broadly, this approach can reveal how different constraints and contexts are integrated with perceptual representations to support effective information processing.

## Appendix

## Computing model gradients

The log-likelihood in Equation 2 is a function of the predicted choice probabilities in Equation 1. To compute the gradient of the log-likelihood, we therefore take the derivative of Equation 2 w.r.t. a given stimulus along dimension *a, s*_*xa*_. Using the chain rule:

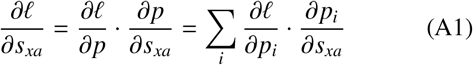

where

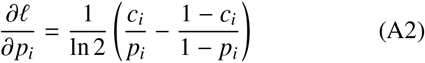

To compute the second part of the gradient, we note that the derivative of the Euclidean distance function, *d*(*s*_*x*_, *s*_*y*_) along stimulus dimension *a* is:

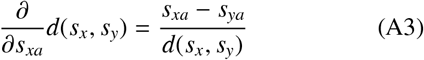

and the derivative of the standard normal PDF, Φ, is the standard normal CDF, ϕ. Because *p*_*i*_ varies as a function of each of the stimuli presented in the triad, the derivative depends on whether *s*_*x*_ was the left, right, or reference stimulus. Therefore, for each trial, we have three possibilities:

1. *s*_*x*_ is the left stimulus, such that

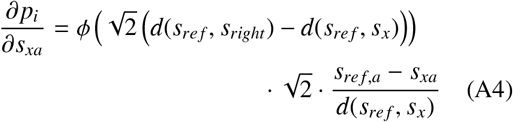
2. *s*_*x*_ is the right stimulus, such that

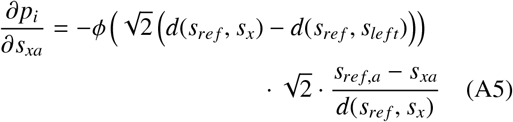
3. *s*_*x*_ is the reference stimulus, such that

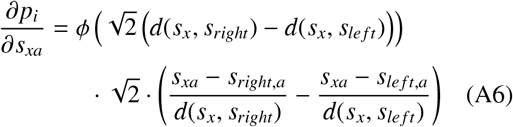

On trials in which *s*_*x*_ was not shown, the gradient is zero. We can then enter the derivatives from each trial into Equation A1 to obtain the overall gradient given the input coordinates.

**Figure S1.**
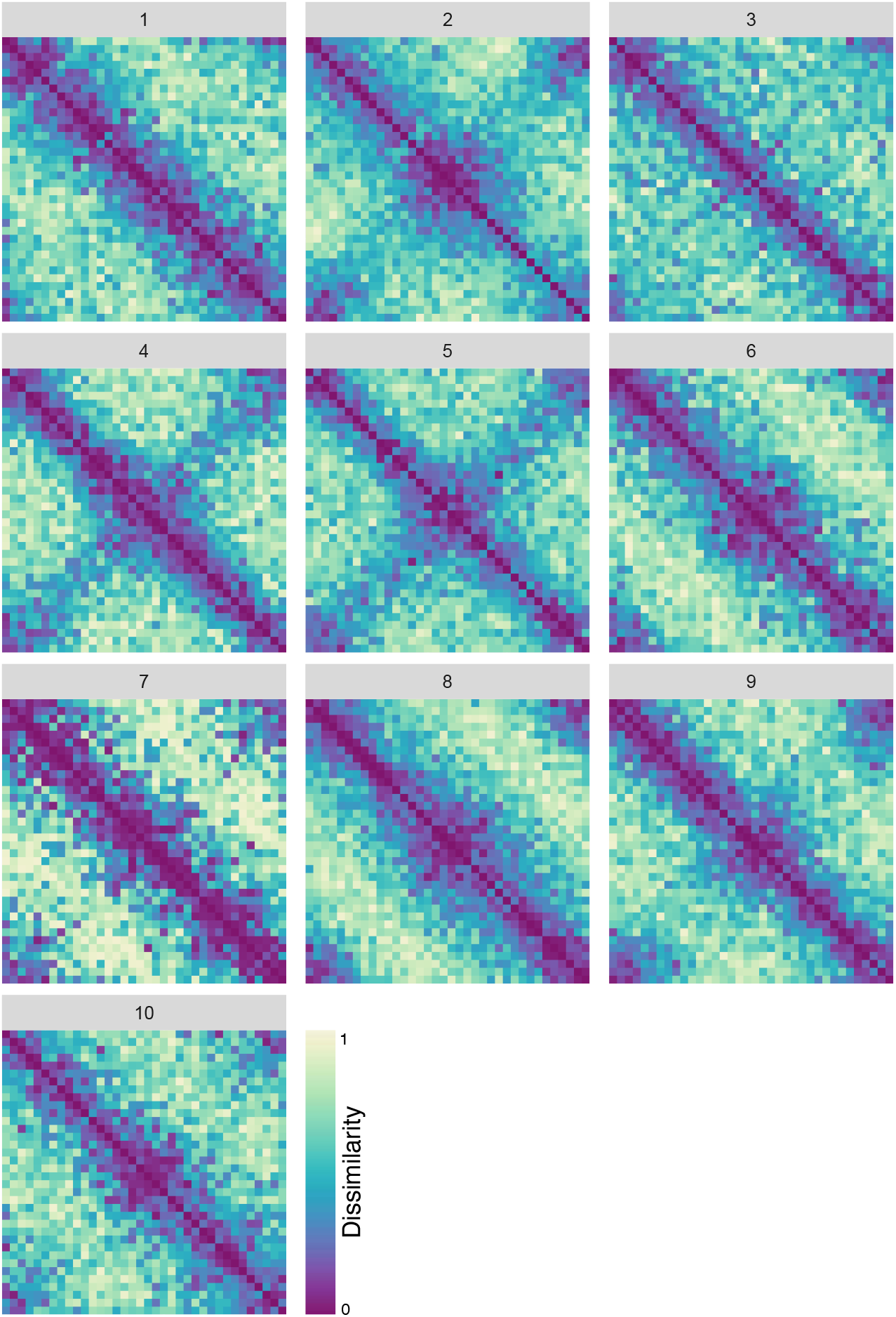
Individual representational dissimilarity matrices across all trials in the similarity judgment task. Differences across participants’ similarity judgments could be driven by variation in perception or decision processes

**Figure S2.**
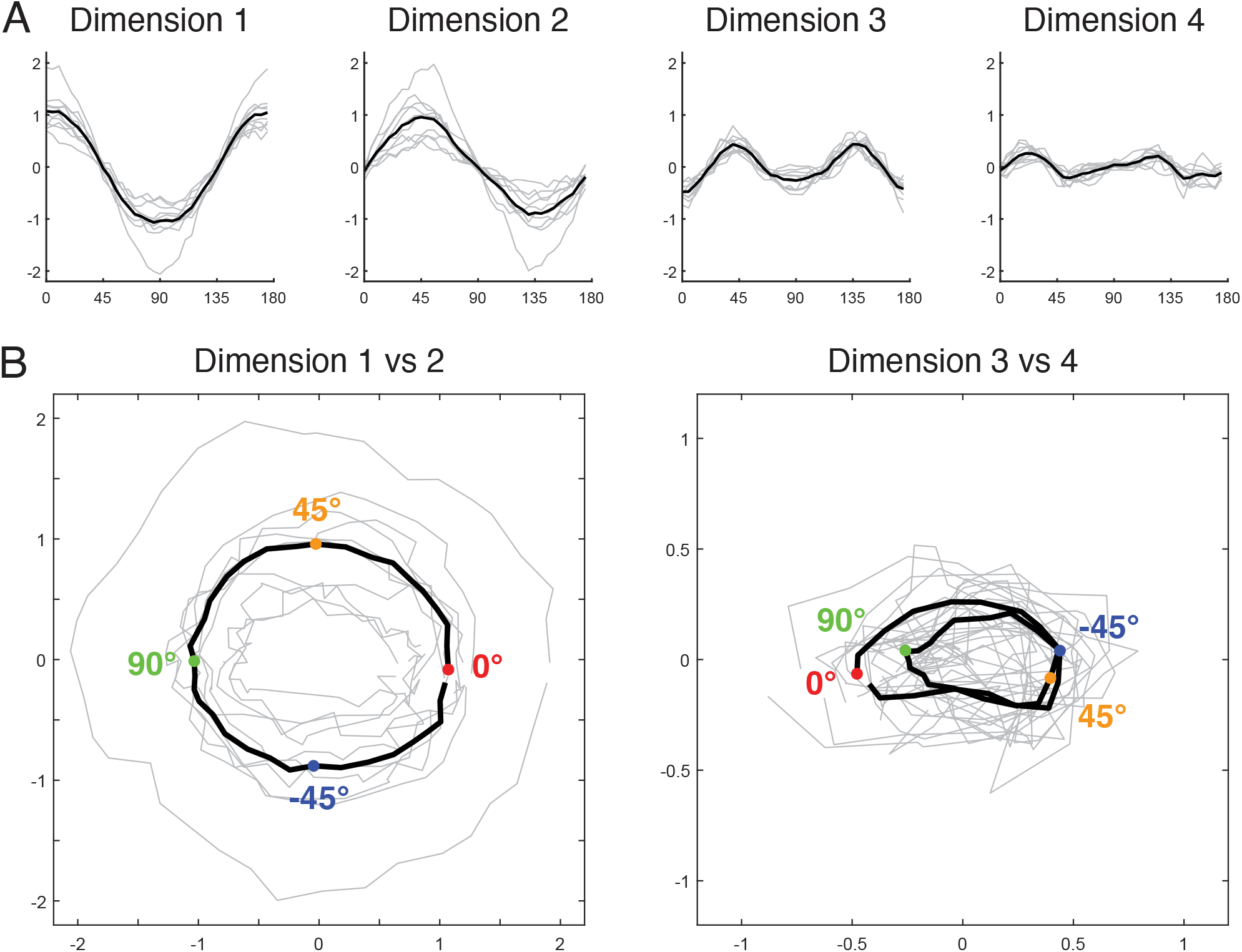
A) Solution of the geometric behavioral model in 4-dimensions. Each dimension is plotted separately as a function of stimulus orientation. Light grey lines represent individual participant’s coordinates, while black lines show the group average. B) When plotted against one another, the first two dimensions show a circular structure, while the second two dimensions show a “double loop”.

**Figure S3.**
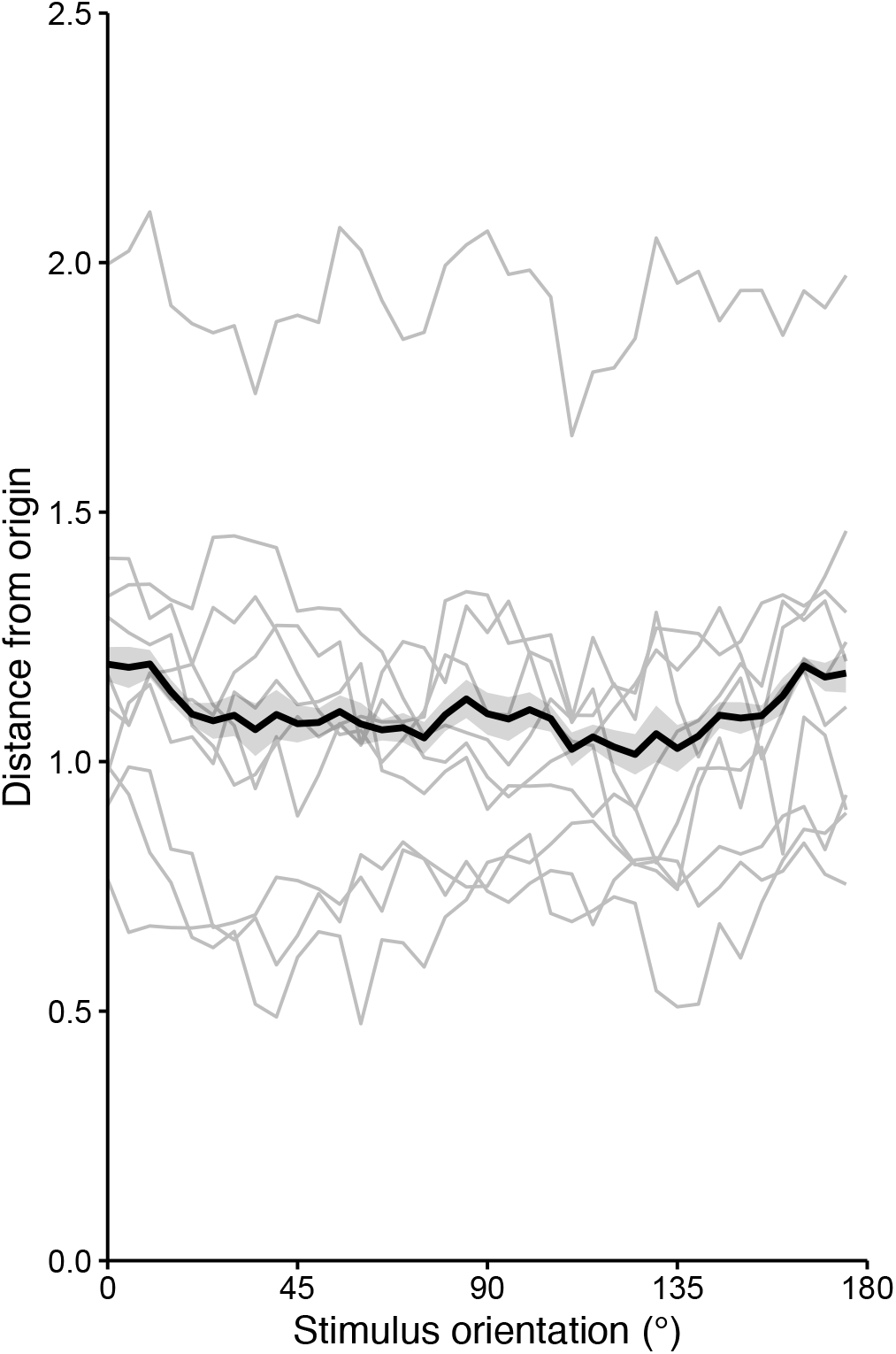
Euclidean distance of each stimulus from the origin using the 4-D model solution. Overall, cardinal orientations (0°/90°) were further from the origin than oblique orientations (±45°). Light grey lines represent individual participants, while the black line shows the group average. Shaded regions correspond to within-subjects SE.

**Figure S4.**
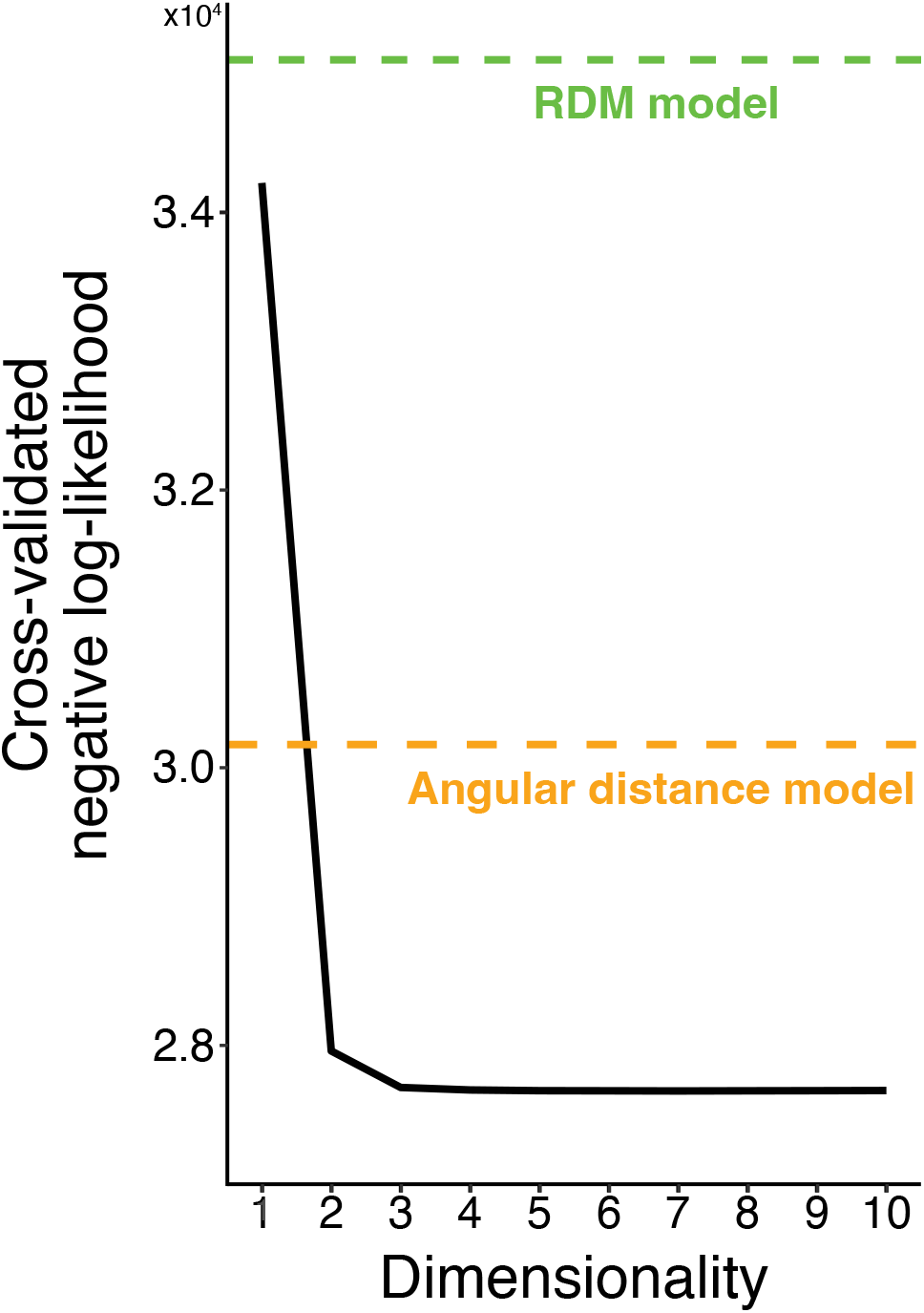
Cross-validated negative log-likelihood of the best fitting behavioral model at each dimensionality, with coordinates that were allowed to vary across attention conditions. Model fits improved up to around 3-4 dimensions, at which point the negative log-likelihood remained nearly constant. Horizontal dashed lines indicate the fit of two the angular distance model and the RDM model.

**Figure S5.**
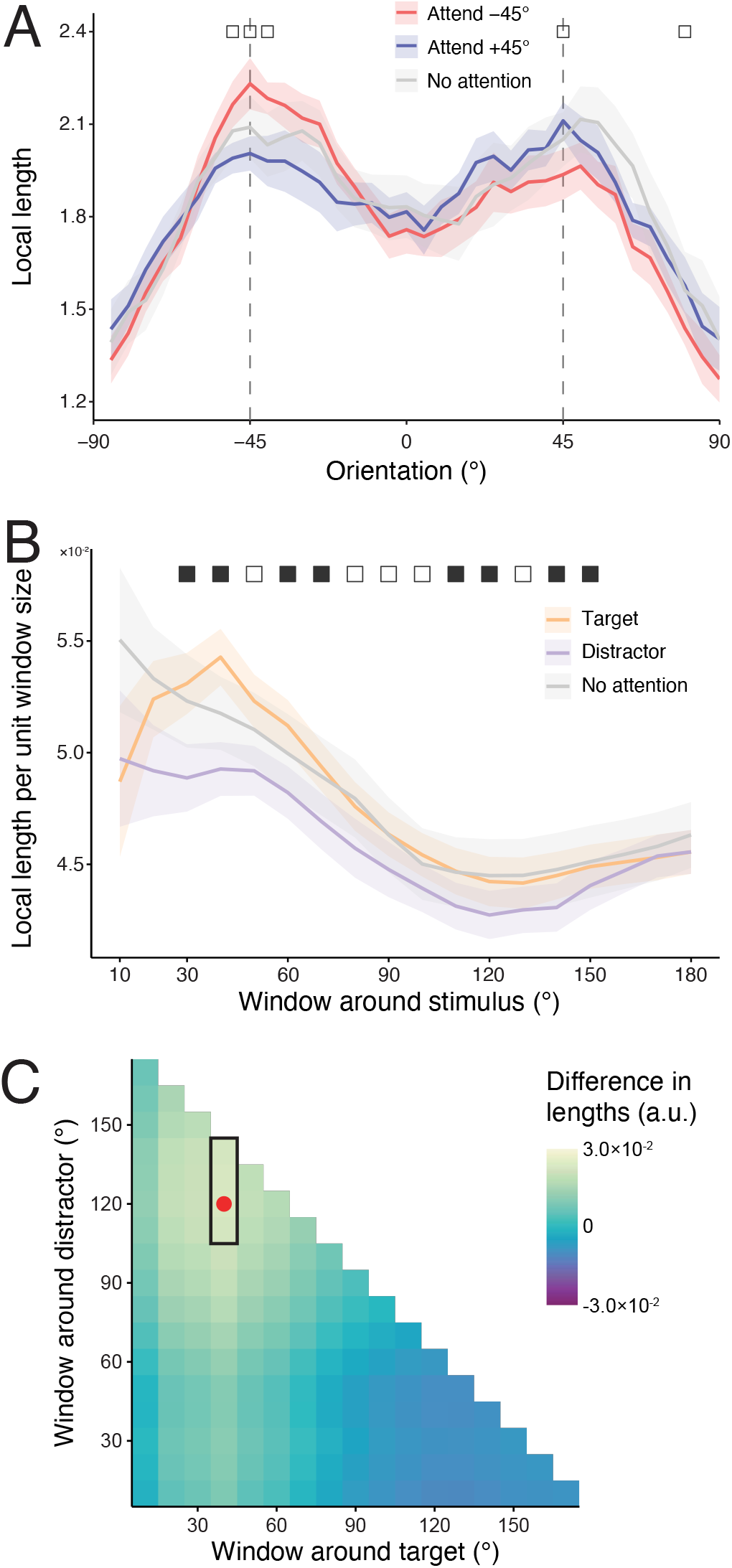
Expansion and compression of orientation representations using the original (non-normalized) lengths. A) Local lengths of perceptual representations of orientations in each attention condition. B) Local length around ±45° orientations by window size when it was the target or distractor orientation, or when no orientation was attended. C) Window sizes maximizing relative expansion around targets vs. compression around distractors. All conventions follow Figure 4.

**Figure S6.**
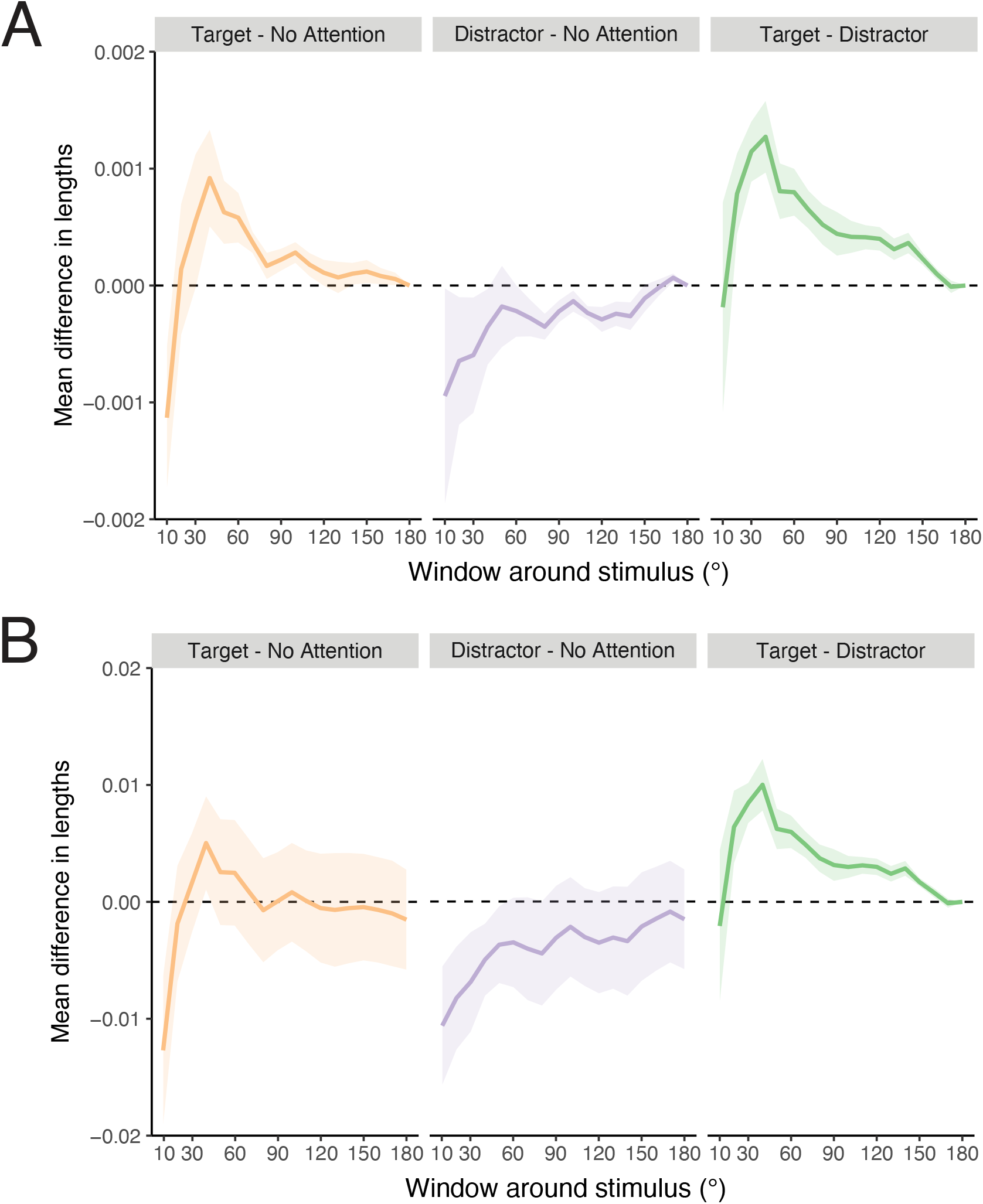
Comparisons of mean local length between different attention conditions, for the A) normalized, and B) original lengths. For a given window size, we calculated the mean local length for each condition and then calculated the difference between each pair of conditions. Shaded regions correspond to SE of the difference.

**Figure S7.**
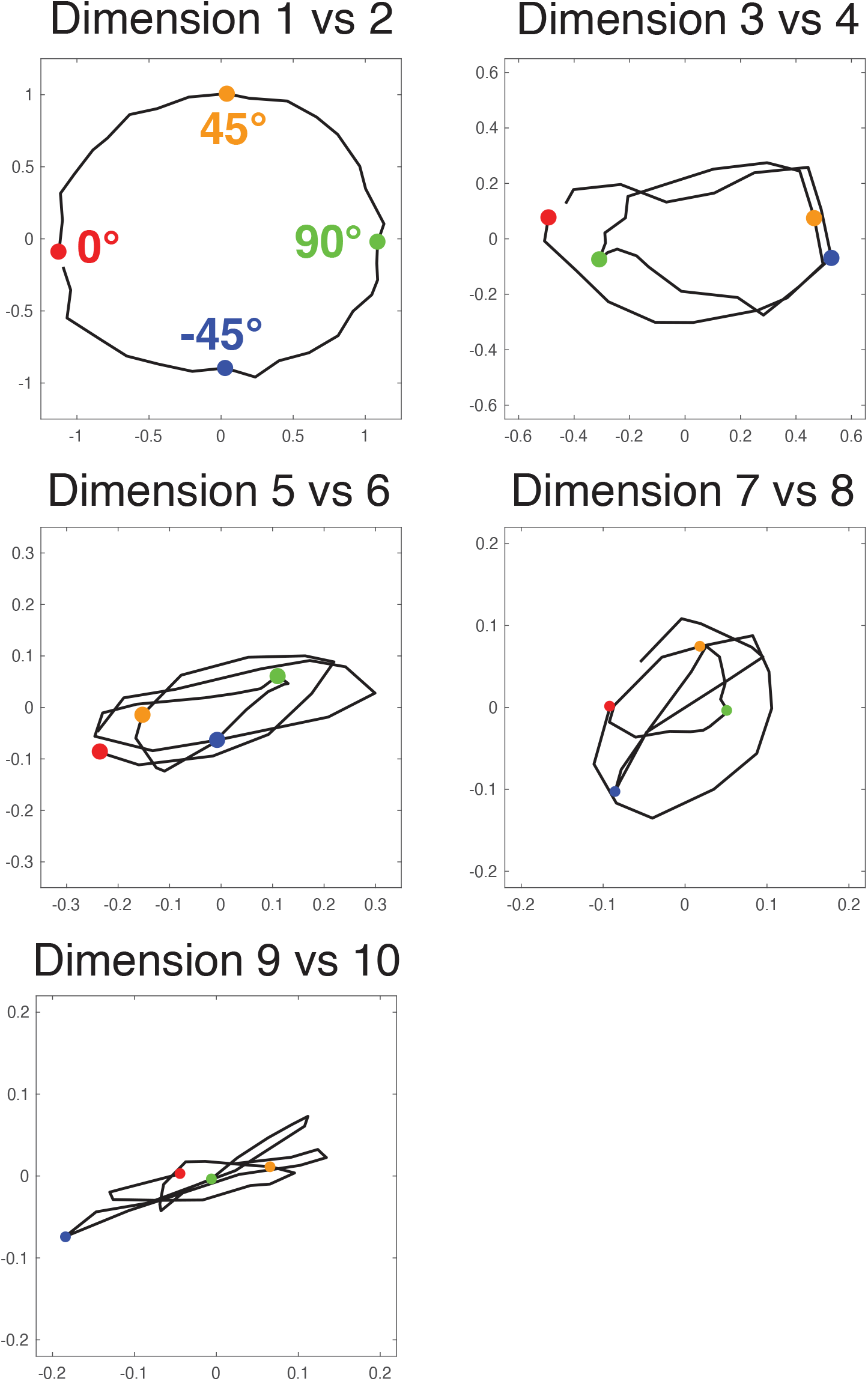
Loops in the 10-dimensional geometric model solution, plotted for pairs of successive dimensions. The first three pairs show a single-, double-, and triple-loop respectively, while the last two pairs do not show a clear increase in the looping structure.

